# Burst suppression: a default brain state associated with loss of network complexity

**DOI:** 10.1101/2025.10.21.683673

**Authors:** Nina Doorn, Gerco C. Hassink, Monica Frega, Michel J.A.M. van Putten

## Abstract

Burst suppression (BS) is a highly stereotyped EEG pattern observed across a wide range of clinical contexts, from general anesthesia and postanoxic coma to neonatal encephalopathy. Despite its consistent appearance, BS comprises two distinct forms with markedly different implications. BS with identical bursts (IBS) is almost exclusively seen in patients with severe, irreversible encephalopathy and is consistently associated with poor neurological outcome. In contrast, heterogeneous BS (HBS) can appear in reversible conditions such as anesthesia. The mechanisms that give rise to these divergent forms remain elusive. Existing theories impose disease-specific processes on otherwise healthy networks, but such models fail to explain why BS emerges across diverse etiologies and disregard the clinically critical distinction between IBS and HBS.

We combined clinical, experimental, and computational approaches to identify shared mechanisms underlying BS. We analyzed EEG recordings from patients with a severe postanoxic encephalopathy and from patients undergoing general anesthesia. These clinical observations were compared with activity recordings from human induced pluripotent stem cell-derived neuronal networks and rodent cortical cultures, and simulations of biophysically grounded neuronal network models.

Purely excitatory, low-complexity networks, both *in vitro* and *in silico*, spontaneously generated activity virtually indistinguishable from pathological IBS. Introducing inhibitory neurons, modular network structure, or diverse external inputs progressively increased signal complexity and produced HBS-like or continuous activity resembling physiological EEG.

Our findings suggest that BS, and particularly IBS, reflects a default dynamic state of simplified excitatory networks that emerges when biological complexity is lost. Different clinical conditions may compromise distinct mechanisms—inhibition, connectivity, or afferent input—yet converge on the same underlying activity pattern. While IBS reflects near-complete loss of complexity, HBS indicates partial preservation. This unified framework explains how diverse etiologies converge on BS and highlights identical forms as signatures of severely reduced network complexity.

## Introduction

Burst suppression (BS) is a distinctive EEG pattern characterized by alternating high-amplitude bursts and periods of suppression. It remains one of the most enigmatic patterns in neurology, appearing across vastly different clinical conditions, from general anesthesia to postanoxic coma to neonatal encephalopathies^1–4^. This raises a fundamental question: how can such diverse pathologies produce nearly identical brain activity patterns?

BS patterns vary in their characteristics and prognostic significance^1^. Heterogeneous burst suppression (HBS) shows variable burst morphology and can be observed in both pathological and physiological situations. HBS can be induced with general anesthetics^5,6^, where the duration of suppression reflects anesthetic depth^7–9^. HBS can also be medically induced for its potential neuroprotective effect in traumatic brain injury, refractory status epilepticus, and cardiac surgery^10–13^. Additionally, HBS arises physiologically in preterm infants and during normal neonatal sleep (tracé alternant)^14^. In contrast, identical burst suppression (IBS) consists of highly stereotyped discharges with abrupt onsets and near-complete suppression between bursts^15^. IBS appears in postanoxic coma^16,17^, severe neonatal asphyxia^18^, and early infantile epileptic encephalopathies^19^, and invariably predicts poor neurological outcome^16^.

Existing theories cannot explain this convergence of diverse etiologies on BS, nor account for the variance of BS patterns and their diverging prognostic implications. The metabolic depletion hypothesis^20^ proposes that ATP deficiency drives suppression through activation of ATP-gated potassium channels. However, this model produces stereotyped pure *α*-band bursts not observed clinically, cannot distinguish IBS from HBS, and struggles to explain BS during anesthesia (where ATP demand is reduced^21^). The cortical hypersensitivity hypothesis^22,23^ suggests that during BS, the cortex becomes hyperexcitable, with minimal stimuli triggering global bursts. While this describes the phenomenon well, it does not explain what causes this hypersensitivity across different pathologies, what drives the suppression phases, or what differentiates HBS from IBS. Other theories suggest that BS may be driven by subcortical pacemakers such as the thalamus or brainstem, based on evidence from EEG source modeling^24,25^. However, BS has also been recorded in isolated cortical preparations without subcortical input^26,27^, suggesting that an intrinsic mechanism of cortical networks can generate BS. Finally, generative models show BS requires a fast-slow dynamical system^3,17^. While this may capture the core dynamics, these models typically impose biologically questionable slow modulators onto otherwise healthy networks, and they do not account for the full phenomenology of BS: its convergence across pathologies, the distinction between IBS and HBS, and its prognostic divergence.

Here we propose a paradigm shift: rather than asking what creates BS, we ask what normally prevents it. We hypothesize that BS, particularly IBS, represents the default activity state of minimal excitatory neuronal networks, which is revealed when healthy biological complexity fails. This perspective is motivated by the observation that the simplest *in vitro* networks, consisting only of randomly connected excitatory neurons, spontaneously produce IBS-like activity. These minimal cultures thus reveal the intrinsic tendency of excitatory networks toward BS. In our experiments and biophysical simulations, the addition of complexity—through inhibition, structured connectivity, and diverse inputs—consistently reduced bursting and increased signal complexity. We propose that, in patients, BS emerges when one or more of these complexity-generating mechanisms are compromised; different etiologies act via different routes yet converge on the same default pattern. This framework explains both why BS appears across diverse clinical conditions and why IBS consistently signals poor prognosis: it represents the most severe loss of complexity.

## Results

### Simple excitatory networks spontaneously generate identical burst patterns

Clinical EEG recordings reveal a spectrum of BS patterns. Under anesthesia, BS emerges with heterogeneous bursts that vary in shape and timing (Figure 1B). Postanoxic coma patients show either heterogeneous burst suppression (HBS) characterized by gradual transitions between bursts and suppressions (Figure 1C,F), or identical burst suppression (IBS): highly stereotyped discharges with abrupt onsets and a nearly completely suppressed EEG (*<* 10 µV) between the bursts (Figure 1D,F). In contrast, awake individuals show continuous *α*-dominated activity (Figure 1A).

**Figure 1.**
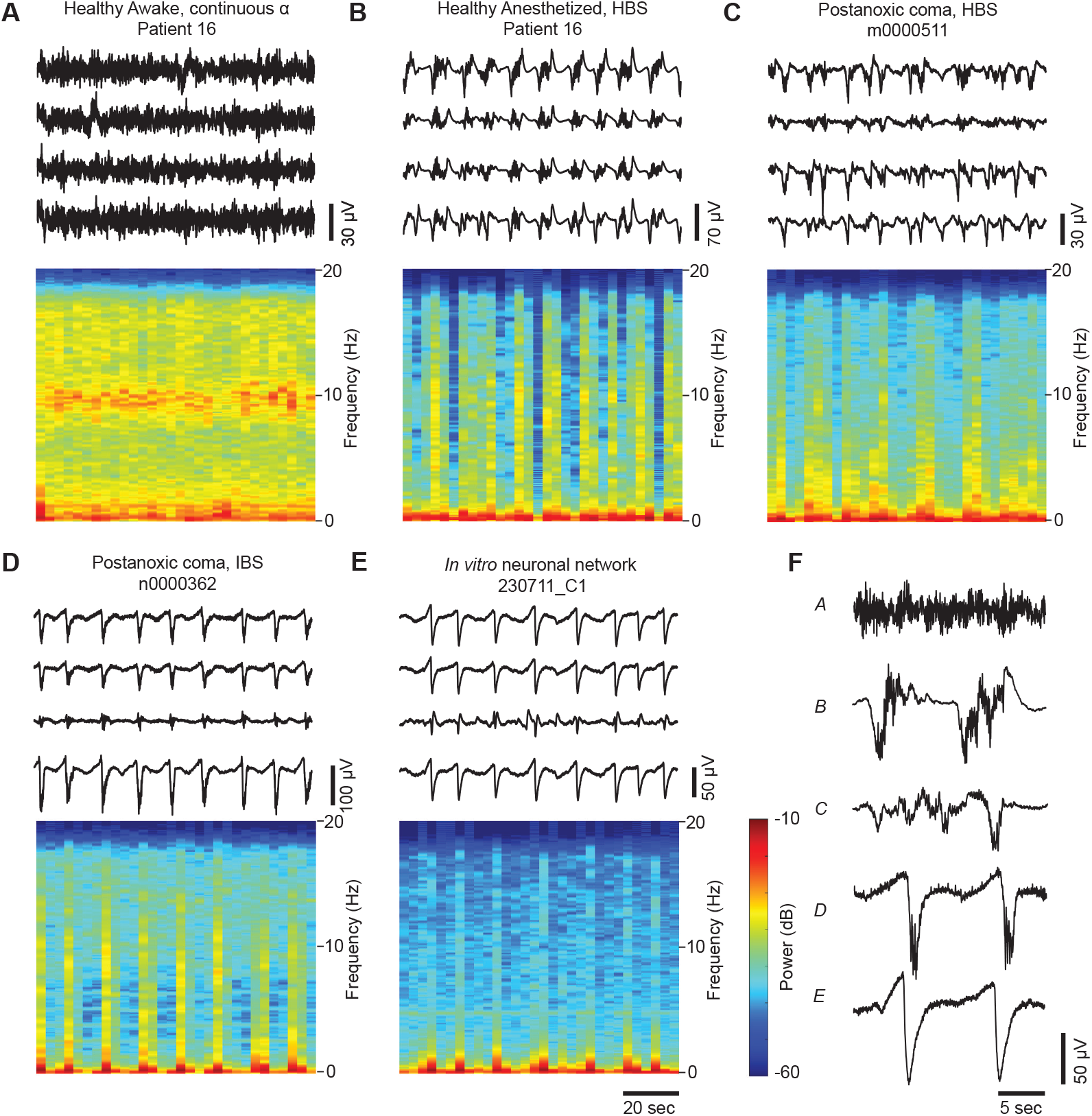
Burst suppression patterns vary across etiologies, with simple excitatory cultures matching the most pathological form. **A-D)** Top: 100 s of EEG (filtered 0.1-25 Hz) from four representative leads and bottom: average spectrogram from all EEG leads from **A)** an otherwise healthy subject with eyes closed before undergoing carotid surgery, the EEG mostly shows continuous *α* activity (at around 10 Hz), **B)** the same patient after propofol application, the EEG shows burst suppression (BS) with heterogeneous bursts (HBS), **C)** a patient in coma after cardiac arrest, EEG showing HBS, **D)** a patient in coma after cardiac arrest, EEG showing BS with identical bursts (IBS). **E)** Top: A 100 s multi-electrode array (MEA) recording (filtered 0.1-25 Hz) of the spontaneous activity of an *in vitro* human induced pluripotent stem cell (hiPSC)-derived excitatory neuronal network. Four representative electrodes are shown. Bottom: average spectrogram of all MEA electrodes. **F)** Representative 20 s epochs from one lead/electrode of the respective panels A-E. Note the resemblance between *in vitro* bursts and identical EEG bursts.

Notably, *in vitro* excitatory neuronal cultures spontaneously generate activity virtually indistinguishable from pathological IBS (Figure 1E). These cultures, which we will refer to as “simple” networks, are comprised of a couple thousand randomly connected excitatory neurons in 2D, differentiated from human induced pluripotent stem cells (hiPSCs) and recorded using multi-electrode arrays (MEAs)^28^. These networks naturally produce BS patterns with abrupt burst onsets, dominant rhythms in the *δ*-band during bursts, and complete inter-burst suppression, closely mirroring the most severe clinical cases.

### Complexity analysis quantifies simple cultures at the pathological extreme of the healthy-IBS spectrum

To systematically characterize differences across the clinical spectrum, we quantified two key features of brain activity: Lempel-Ziv complexity, reflecting pattern predictability, and burstiness (coefficient of variation of voltage changes), capturing temporal intermittency. These metrics showed a clear gradient across the clinical spectrum. Healthy awake EEG exhibited maximum complexity and minimum burstiness, consistent with continuous, variable activity (Figure 2A). Anesthesia-induced HBS showed intermediate values, while pathological IBS dropped to minimal complexity and maximal burstiness. Crucially, excitatory cultures displayed complexity values indistinguishable from IBS EEG, quantitatively confirming that these simple networks reproduce the most pathological brain activity patterns.

**Figure 2.**
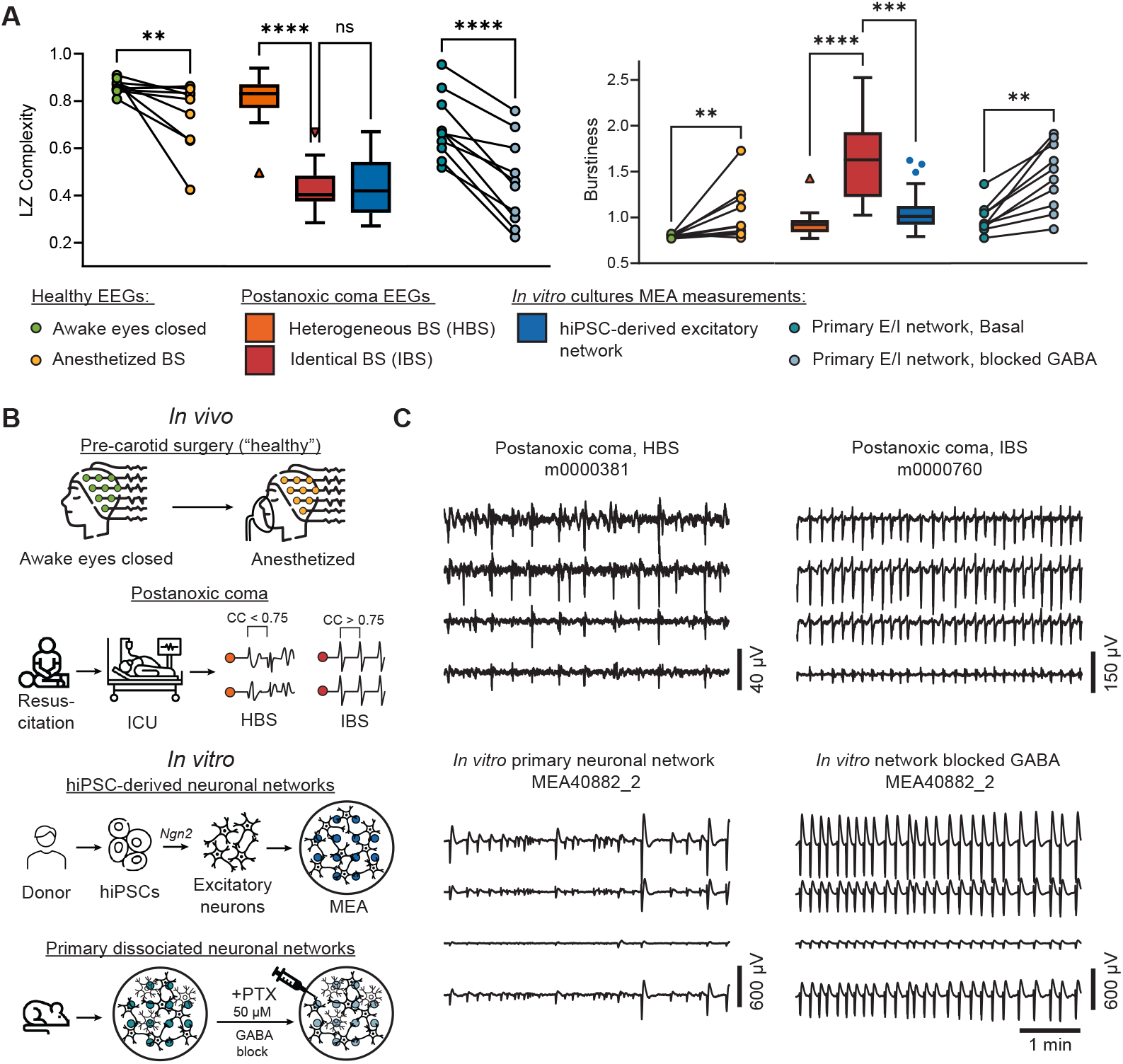
Blocking inhibition in mixed cultures recreates the transition from HBS-like to IBS-like patterns. **A)** Quantification of the Lempel-Ziv (LZ) complexity and the burstiness (coefficient of variation of the absolute value of the derivative) of the EEGs from patients before undergoing carotid surgery (“healthy”), before and during anesthesia (*n* = 10), from patients in postanoxic coma showing heterogeneous burst suppression (HBS) (*n* = 18) and identical BS (IBS) (*n* = 15), of the MEA recordings from human induced pluripotent stem cell (hiPSC)-derived excitatory neuronal networks (*n* = 29) and from primary rodent neuronal cultures with excitatory and inhibitory neurons, before and after the addition of 50 µM PTX to block GABA receptors (*n* = 10). Paired measurements (Awake vs. Anesthetized and Basal vs. blocked) are compared using a Wilcoxon matched-pairs signed rank test. Non-paired measurements are compared using a Kruskal-Wallis test with Dunn’s multiple comparisons test. ns *p >* 0.05, ** *p <* 0.01, *** *p <* 0.001, **** *p <* 0.0001. Note the observations in the plots are compared with multiple independent statistical tests. **B)** Schematic overview of how the data quantified in panel A is obtained. **C)** Top: representative example 5-minute EEG recordings from postanoxic coma patients showing heterogeneous and identical BS (HBS and IBS). Bottom: representative example 5-minute MEA recording from a rodent neuronal culture before and after blocking GABA receptors.

Having placed excitatory cultures at the IBS extreme, we next turn to them as a controlled testbed. We build on extensive culture studies showing that inhibition, structured connectivity, and external input are key determinants of network dynamics^29–34^. Here, we use the *in vitro* system and matched *in silico* models to quantify how these properties shift signals along the LZ-complexity and burstiness axes relative to IBS, HBS, and healthy EEG. By delineating their effects on these metrics, we test the default-state hypothesis: simple networks naturally express IBS, whereas adding complexity suppresses bursting and steers activity toward healthy-like, continuous dynamics.

### Inhibition shifts cultures off the IBS extreme

To probe the role of inhibitory neurons, we first examined mixed excitatory–inhibitory rodent cortical cultures, which contain a diverse pool of cortical cell types and typically show richer dynamics than our hiPSC-derived excitatory-only cultures^35^. Our mixed cultures exhibited HBS-like activity with higher complexity and lower burstiness (Figure 2A,B).

To isolate the effect of inhibition from the effect of species differences and increased cellular heterogeneity, we then acutely blocked inhibitory neurotransmission in the *same* mixed cultures using the GABA_A_ antagonist picrotoxin. This single manipulation transformed their signals from HBS-like to IBS-like: LZ complexity dropped and burstiness increased dramatically (Figure 2A,C), matching both our excitatory-only cultures and clinical IBS. Thus, when species and cell composition are held constant, removing inhibition alone is sufficient to drive cultures to the IBS extreme.

### Network structure restores complexity and burstiness under low inhibition

Beyond inhibition, prior work shows that more structured culture architectures yield richer dynamics^30–32,36^. Our hiPSC-derived networks approximate random connectivity. Motivated by this literature, we asked how introducing structured connectivity would interact with inhibitory tone to shift LZ complexity and burstiness.

Using our validated biophysical model of hiPSC-derived networks^37,38^, we systematically explored the interaction between inhibition and modular architecture. We incorporated inhibitory neurons and implemented a modular structure with parameter Q controlling the ratio of intra-module to inter-module connections (Figure 3B). Our model with a modular structure mimicked the activity of modular cultures of Yamamoto et al.^36^ (Supplementary Figure S1). Moreover, although we only fitted the model to high-frequency activity features of hiPSC-derived neuronal networks (action potential-related features), the low-frequency component of the simulated signals mimicked that of EEG BS recordings and experimental MEA recordings (Supplementary Figure S2), with dominant power in the *δ* band.

**Figure 3.**
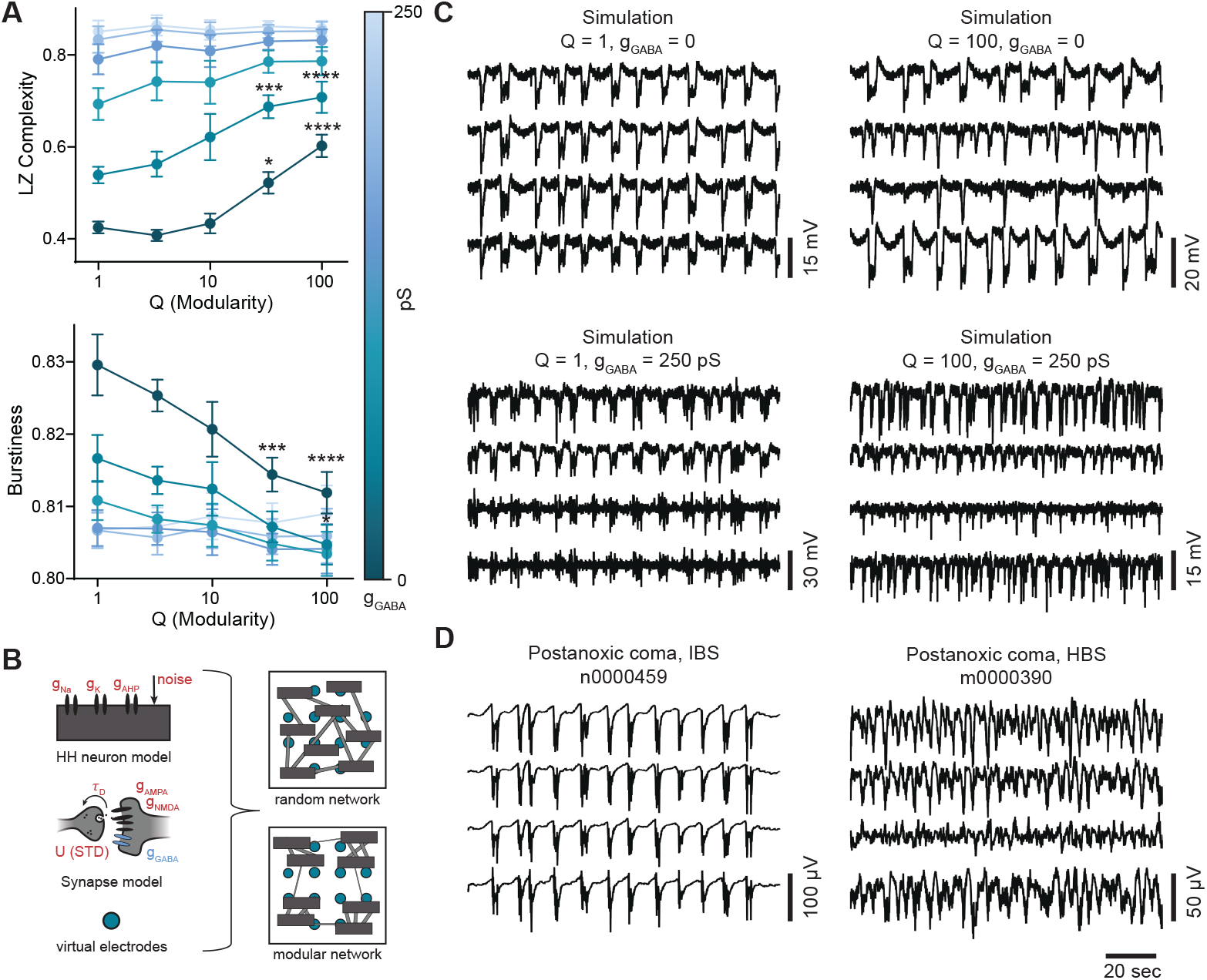
Network structure rescues activity from IBS-like patterns only when inhibition is limited. **A)** Quantification of the Lempel-Ziv (LZ) complexity and burstiness for computational model simulations with increasing network modularity (Q) for six equally spaced values of the maximum GABA receptor conductance (*g*_GABA_). Only with low *g*_GABA_, modular networks show higher complexity and lower burstiness than random networks. Plots show mean±SEM, *n* = 10 per condition (one value of *g*_GABA_ and one value of Q). Per *g*_GABA_ value, quantifications with Q*>*1 are compared to the quantification of Q=1 (a random network) using a 2-way ANOVA and Dunnett’s multiple comparisons test. *** *p <* 0.001, **** *p <* 0.0001, if not indicated *p >* 0.05. **B)** Schematic overview of the computational model consisting of 100 excitatory and 40 inhibitory Hodgkin-Huxley (HH) type neurons and conductance based synapses. The parameters in red are fit using simulation-based inference. **C)** Representative 100 s example simulations with different values for *g*_GABA_ and Q. Networks with low Q and *g*_GABA_ resemble highly pathological identical burst suppression (IBS), while increasing *g*_GABA_ or Q increases signal complexity, resembling heterogeneous BS (HBS). **D)** Representative example 100 s EEG recordings from postanoxic coma patients showing HBS and IBS.

### Diverse inputs destabilize stereotyped bursting

Synchronized bursting in neuronal cultures has been linked to the absence of diverse afferent inputs that normally desynchronize activity *in vivo*^34,39^. *Moreover, distributed electrical stimulation has been shown to reduce burstiness*^*34*^.

We modeled this by increasing stochastic membrane potential fluctuations (input “noise”) in our computational framework to mimic diverse afferent drive. To maintain comparable activity levels, we proportionally reduced neuronal excitability as noise amplitude increased. Increasing noise amplitude drove networks away from stereotyped toward variable bursting patterns, elevating complexity and reducing burstiness (Figure 4A). With minimal noise, constant-current inputs generated highly predictable, IBS-resembling bursts. Progressive noise increases diversified burst morphologies, producing more variable, HBS-like patterns (Figure 4B).

**Figure 4.**
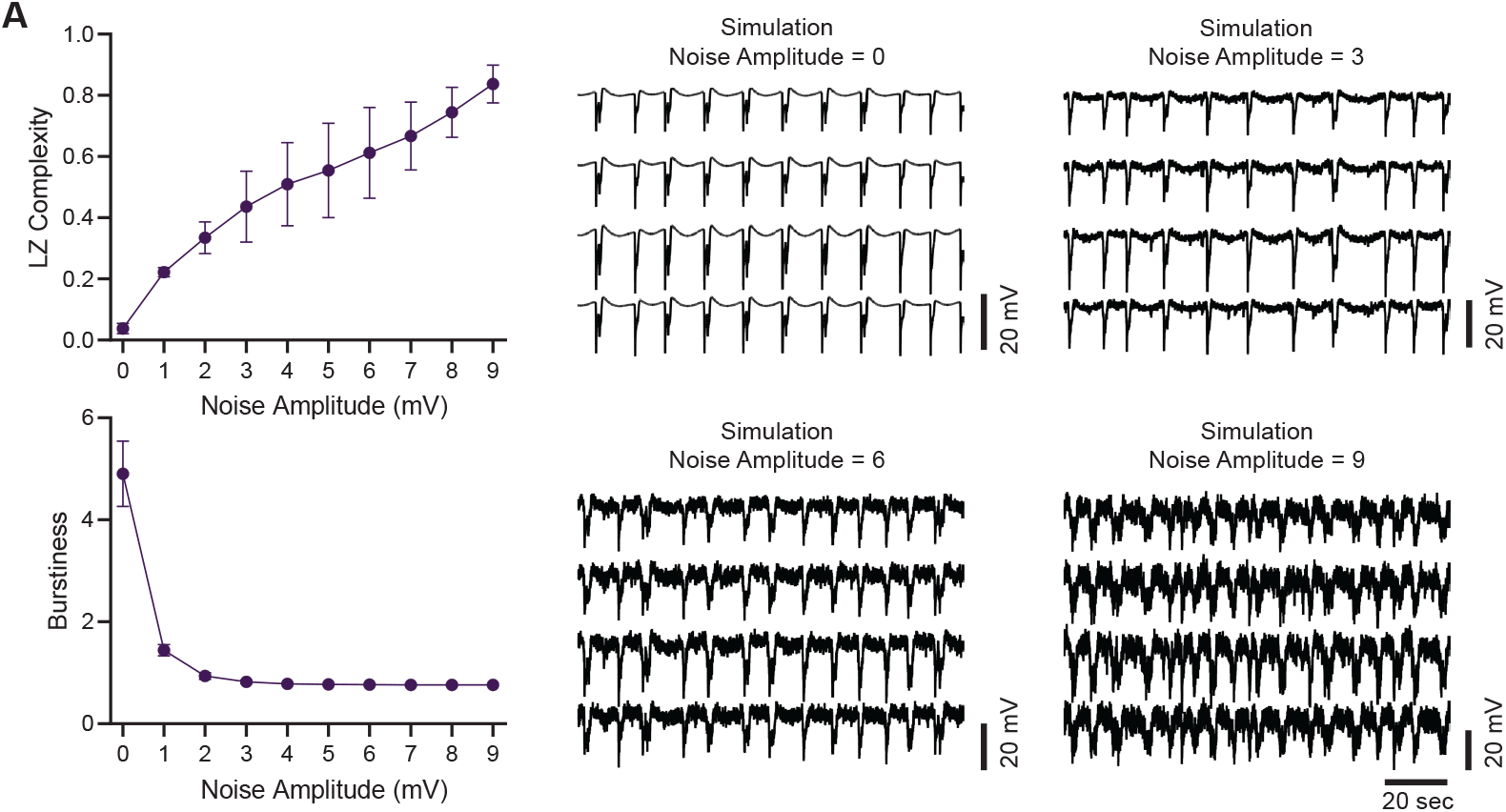
Computational modeling shows diverse inputs can drive simulated networks from IBS-like to HBS-like patterns. Quantification of the Lempel-Ziv (LZ) complexity and burstiness for model simulations with different amplitudes of the noisy membrane potential fluctuations. Plots show mean ± SEM, *n* = 10 per condition. **B)** Representative 100-second example simulations with increasing values of the noise amplitude. Higher noise amplitude values result in simulations showing more heterogeneous bursts with higher complexity. Completely random networks (Q = 1) required substantial inhibition to achieve complexity in-creases and burstiness reductions. However, structured networks (higher Q) achieved similar effects, but only with no or minimal inhibition (Figure 3A). Networks with low inhibition and low modularity generated stereotyped, IBS-like patterns, while increasing either inhibition or modularity produced more variable, HBS-like dynamics (Figure 3C,D).

*Taken together, our in vitro* cultures and matched *in silico* models indicate that complex, heterogeneous activity is supported by three factors: the presence of functional inhibition, structured (e.g., modular) connectivity, and diverse inputs. When the system is simplified—excitatory-only, random 2D connectivity, and/or deprived of input—IBS-like stereotyped bursting emerges. These convergent effects suggest plausible routes to BS-like patterns via reduced inhibition, disrupted structure, or diminished inputs, which we consider further in the Discussion.

## Discussion

We identify identical burst suppression (IBS) as a default dynamic of purely excitatory networks, one that emerges spontaneously when key regulatory mechanisms are absent. Using neuronal cultures, clinical EEG, and computational models, we show that IBS arises naturally in simple networks lacking three critical forms of regulation: synaptic inhibition, structured connectivity, and diverse inputs. In contrast, heterogeneous burst suppression (HBS) requires at least partial preservation of one or more of these mechanisms. This fundamentally reframes how we approach BS: rather than being caused by specific disease processes, IBS emerges when the systems that normally prevent pathological synchrony break down.

This paradigm shift explains a puzzling clinical observation: why does a similar EEG pattern appear across vastly different conditions, from anesthesia to cardiac arrest to neonatal encephalopathy? Current computational models struggle with this diversity. The prominent theory by Ching et al.^20^ proposes that BS is caused by reduced ATP production leading to activation of ATP-gated potassium channels, temporarily silencing neuronal activity until ATP levels are restored. The authors claim that all BS etiologies can be linked to reduced rates of ATP production, directly in anoxia and indirectly in anesthesia through down-regulation of neuronal firing by the anesthetic and a subsequent autoregulatory decrease in cerebral metabolic rate of oxygen. However, in the latter case, ATP consumption would also be reduced, making a net ATP deficit unlikely. In the postanoxic coma case, BS is often observed long after circulation, and presumably ATP production, has been restored, challenging the ATP-hypothesis. Liley and Walsh^3^ showed that adding short-term synaptic depression to a mean-field model of cortical activity can produce BS-like alternations. Both the ATP-depletion model of Ching et al. and the synaptic-depression model of Liley and Walsh generate artificial *α*-band bursts never seen clinically, and they ignore the fundamental distinction between HBS and IBS, despite its major prognostic significance. Moreover, both approaches add complexity (ATP depletion or synaptic depression) to otherwise healthy brain models to force BS, and thus fail to explain how diverse clinical conditions converge on BS.

Our approach reverses this logic. We demonstrate that the most pathological EEG pattern, IBS, arises naturally from the activity of a minimal neuronal architecture: a randomly connected 2D excitatory network. In contrast, more variable patterns, such as HBS or physiological activity, require the presence of regulatory mechanisms that prevent this stereotyped default activity.

### The cellular machinery underlying the default state

What drives this default bursting pattern? In our computational models, BS emerges from a classic fast-slow dynamical system: rapid recurrent excitation (fast) coupled with slow adaptation currents (slow)^38^. When neurons spontaneously fire together due to external input or intrinsic noise, positive feedback through excitatory connections triggers network-wide bursts. This intense activity gradually activates slow afterhyperpolarizing potassium currents, eventually silencing the network until this adaptation sufficiently recovers and the cycle repeats. Previous models revealed fast-slow dynamics can generate BS, but they invoked implausible slow modulators like synaptic depression onto otherwise healthy networks to transition to BS^3,17^. Our framework inverts this reasoning: the adaptation-based fast-slow system exists intrinsically in all excitatory networks, but complexity-generating mechanisms suppress its BS manifestation in healthy networks.

This adaptation-based mechanism aligns with extensive literature showing that adaptation currents underlie bursting patterns and slow cortical oscillations across diverse neuronal systems^40–44^. The adaptation framework also explains key experimental observations. Kroeger and Amzica measured hyperpolarized membrane potentials during the suppression phases of BS, exactly what strong adaptation currents would produce^22^. They also documented a characteristic refractory period: immediately after a burst, when adaptation currents are strongest, external stimuli cannot trigger another burst. However, as these currents slowly decay over time, stimuli become progressively more effective at eliciting bursts. The absolute refractory period lasted 2.5 seconds, matching the minimal decay time constant of slow afterhyperpolarizing currents^45,46^ but an order of magnitude larger than typical synaptic depression timescales^47^. Remarkably, this exact refractory phenomenon also occurs in postanoxic coma patients, where minimal stimuli can trigger whole-brain bursts, but only after sufficient time has passed since the previous burst^48^. Moreover, the refractory pattern appears in neuronal cultures^49^, suggesting a conserved adaptation-based mechanism operates across scales from cell cultures to human brains.

Together, our findings show that BS can emerge from basic excitation–adaptation dynamics, without requiring complex pathology. This raises a deeper question: if such dynamics are intrinsic to neuronal circuits, what prevents the healthy brain from spontaneously entering this default state? Our experiments reveal three distinct mechanisms that counteract this default bursting.

### Loss of inhibition unveils the default state

We demonstrate that blocking inhibitory neurotransmission transforms variable HBS-like patterns into stereotyped IBS-like activity, suggesting that inhibitory dysfunction plays a central role across BS etiologies.

The clearest case is BS following various forms of anoxic brain injury, which may result from cardiac arrest, stroke, drowning, and perinatal asphyxia^16–18^. Extensive research shows that oxygen deprivation preferentially damages inhibitory neurons and their synaptic connections^50–52^, suggesting inhibition is impaired in patients showing BS after anoxia.

During early development, BS patterns coincide with a critical period when GABA signaling remains excitatory rather than inhibitory due to the postnatal GABA switch^53^. This developmental factor may explain why BS is commonly observed in neonatal sleep and encephalopathies.

Anesthesia presents an apparent paradox: these drugs potentiate GABA receptors yet produce BS. However, Ferron et al.^23^ demonstrated complete suppression of inhibitory potentials during isoflurane anesthesia, with only partial suppression of excitation. Importantly, these effects are dose-dependent: at higher concentrations, anesthetics disproportionately reduce interneuron recruitment, leading to disinhibition^54,55^. This mechanism explains the hyperexcitability Kroeger and Amzica observed^22^, where minimal stimuli could induce whole brain bursts during BS anesthesia but not outside it. Recent studies further support the disinhibition model, showing that anesthesia-induced BS is dominated by pyramidal neuron synchronization while interneurons show no correlation with brain activity^55^. Thus, diverse clinical conditions can converge on BS through a common mechanism: functional loss of inhibition, which unveils the excitatory default state.

### Network structure provides resilience when inhibition fails

Our modeling reveals that organized, modular connectivity can maintain complex activity patterns, especially when inhibition is severely compromised. The influence of structure has direct clinical correlates across BS etiologies.

Patients with a postanoxic encephalopathy show extensive structural connectivity disturbances, with functional network disruptions being most pronounced in those with poor neurological outcomes, the very patients who may show IBS^56,57^. The association between structural damage and pathological BS extends to other conditions, as well: corpus callosum lesions frequently produce BS patterns^58,59^, as does diffuse axonal injury^60^.

Although anesthesia does not cause permanent structural damage, it progressively disrupts the brain’s functional architecture. Studies in anesthetized rats show that specific functional networks gradually disappear under isoflurane, replaced by widespread, spatially non-specific synchronization that resembles the activity of our randomly connected cultures^61^. Similar findings in macaques demonstrate that anesthesia transforms the brain’s modular network structure into a more homogeneous network^62^. This temporary loss of functional modularity may explain why even reversible anesthesia can reveal the same default bursting patterns seen in permanent structural brain damage.

### Subcortical inputs destabilize synchronous bursting

Our simulations reveal how input characteristics shape BS patterns. In our model, intrinsic neuronal noise provides the spontaneous events that trigger burst initiation—without any input, networks primed for BS remain silent. When inputs are sparse but uniform (as is the case in simulations without noise), they generate highly stereotyped bursts. However, when inputs are abundant across neurons, they disrupt the synchrony characteristic of IBS, producing more variable patterns. To isolate this desynchronizing effect from increased burst frequency, we reduced neuronal excitability proportional to noise levels. Note that this may have contributed to the desynchronization. While noise inherently increases LZ complexity, the principle that diverse inputs modulate BS patterns finds strong clinical support.

BS appears during cortical deafferentation, when brain regions lose their normal inputs^63,64^. In healthy brains, abundant and diverse inputs likely prevent BS by disrupting synchronization entirely. During anesthesia, which progressively suppresses sensory responses^65^, reduced inputs may allow BS patterns to emerge. As anesthetic depth increases, fewer inputs remain to trigger bursts, lengthening interburst intervals until activity ceases—explaining why suppression duration reliably tracks anesthetic depth^7–9^.

### Implications and future directions

Our findings offer a mechanistic explanation for why IBS is associated with poor outcomes, while HBS permits recovery. IBS reflects the default state of excitatory networks—highly predictable, low-complexity, and significantly reduced information processing capacity—whereas HBS indicates partial preservation of regulatory mechanisms. This suggests therapeutic strategies might aim to restore inhibition, support network connectivity, or reintroduce input diversity to disrupt pathological synchrony.

While our models successfully transform IBS-like into HBS-like patterns, they cannot generate awake EEG dynamics characterized by continuous *α*-dominated activity, suggesting missing complexity-generating mechanisms. Nevertheless, extensive clinical evidence for inhibitory dysfunction, structural damage, and input deprivation supports our framework for understanding the transition from healthy activity to heterogeneous and identical BS. Rather than a disease-specific mechanism, BS appears to signal a fundamental collapse of the brain’s complexity-generating architecture.

## Materials and Methods

### Patients and EEG recordings

We analyzed EEG recordings from 33 adult patients in coma after cardiac arrest, admitted to the intensive care unit. These patients were previously included in a prospective cohort study examining the predictive value of EEG for neurological outcome^66^. Patients received standard therapy, including hypothermia at 33 °C and sedation with propofol to a level of -4 or -5 on the Richmond Agitation Sedation Scale. Continuous EEG monitoring was performed until patients regained consciousness, died, or for a maximum of five days post-arrest. Two EEG experts identified and classified BS patterns in these recordings^16^. All BS patterns classified as heterogeneous bursts had an average correlation coefficient between burst shapes below 0.75, while identical bursts showed correlation coefficients above 0.75^16^. The Medical Ethical Committee Twente approved the protocol (approval K19-11) and waived the need for informed consent since continuous EEG monitoring is part of standard care in the participating centers.

“Healthy awake” and “Anesthetized BS” EEG recordings were obtained from 10 patients undergoing carotid endarterectomy surgery^67^. EEG monitoring was performed as part of standard clinical care to assess the need for temporary shunting during surgery^68^. Two conditions were analyzed: (1) baseline recordings obtained one day prior to surgery during eyes-closed resting state, from which artifact-free periods were annotated (labeled “Awake eyes closed”), and (2) recordings during propofol anesthesia (200-300 mg/h) displaying HBS, labeled “Anesthetized BS”. All annotations were performed by a board-certified neurologist and clinical neurophysiologist. In all awake patients, the EEG contained a posterior dominant rhythm in the *α*-band (8-13 Hz).

In all EEG recordings, we used 19 Ag/AgCl electrodes placed according to the international 10/20 system. Sampling frequencies ranged from 250 to 1024 Hz, and electrode impedances were kept below 10 kΩ to minimize polarization effects. EEGs were recorded using the NeuroCenter EEG system (Clinical Science Systems, the Netherlands) with a full-band EEG amplifier (TMS International, the Netherlands). A common average reference was used during acquisition; all recordings were subsequently re-referenced to a bipolar montage.

### In vitro cultures and MEA recordings

We cultured hiPSC-derived excitatory neuronal networks on MEAs as previously described^69^. The hiPSC line, reprogrammed from skin fibroblasts of a healthy 30-year-old male, was obtained from the Coriell Institute (GM25256, RRID: CVCL Y8303). Cells were differentiated into excitatory neurons through doxycycline-inducible overexpression of *Neurogenin 2* (*Ngn2*)^28,70^.

Briefly, *Ngn2* -positive hiPSCs were cultured on Geltrex (Thermo Fischer Scientific, #A1413302) in E8 flex medium supplemented with G418 (50 *µ*g/mL) and puromycin (0.5 *µ*g/mL) at 37^*°*^C/5% CO_2_. On day *in vitro* (DIV) 0, cells were plated as single cells onto 24-well MEAs (Multi Channel Systems, Reutlingen, Germany) pre-coated with poly-L-ornithine (50 *µ*g/mL, Sigma Aldrich, #P4957) and human laminin (5 *µ*g/mL, BioLamina, #LN521). Rat astrocytes, obtained from cortices of newborn Wistar rats (P1), were added on DIV 2 in a 1:1 ratio to support neuronal maturation. From DIV 3, medium was switched to Neurobasal medium (Gibco, #21103049) containing DOX (4 *µ*g/mL, Sigma Aldrich, #D9891), B-27 (Thermo Fisher Scientific, #17504044), glutaMAX (2 mM, Thermo Fisher Scientific, #35050061), primocin (0.1 *µ*g/mL, Invivogen, #ant-pm-05), neurotrophin-3 (10 ng/mL, Stemcell, #78074), and brain-derived neurotrophic factor (10 ng/mL, Stemcell, #78005). Cytosine *β*-D-arabinofuranoside (2 *µ*M, Sigma Aldrich, #C1768) was added to eliminate proliferating cells. Medium was changed three times weekly. From DIV 10, 2.5% fetal bovine serum (Sigma Aldrich, #F9665) was added for astrocyte viability. DOX was removed after DIV 14. Cultures were maintained at 37^*°*^C with 80% humidity and 5% CO_2_ until recording on DIV 35-50.

Primary cultures were prepared from isolated cortices of newborn Wistar rats (P1; Janvier Labs) following protocols approved by the Dutch Central Committee on Animal Experiments (CCD) (AVD11000202115663). Cortices were enzymatically dissociated with 0.25% trypsin and triturated. Approximately 80,000 cells were plated onto single-well MEAs (Multi Channel Systems, Reutlingen, Germany) containing 60 titanium nitride electrodes pre-coated with polyethylene imine. Cultures were maintained at 36^*°*^C with 80% humidity and 5% CO_2_ until recording on DIV 21-49. Medium consisted of Neurobasal-glucose-pyruvate (Thermofisher, #15329741) supplemented with D-glucose (1.24 g/L), penicillin-streptomycin-glutamine (100x; Thermofisher, #10378016), B-27 supplement (50x, Thermofisher, #11530536), vitamin C (100x; Sigma Aldrich, #A0278-25G), and nerve growth factor (Sigma Aldrich, #N6009), changed twice weekly.

Spontaneous activity was recorded at a sampling frequency of 10 kHz or 25 kHz for a minimum of 5 minutes while maintaining culture temperature (37^*°*^C for human neurons, 36^*°*^C for rodent neurons) and applying a humidified gas flow (5% CO_2_, 95% air). For pharmacological experiments, we recorded baseline activity for 10 minutes, after which we added 50 *µ*M picrotoxin (PTX, Sigma Aldrich, #P1675), followed by an additional 10 minutes of recording.

### Data processing and analysis

To ensure consistency across recording systems, 10 channels were analyzed. For EEG recordings, frontal and temporal leads (Fp2-F8, F8-F4, T4-T6, Fp1-F7, F7-T3, T3-T5, Fp2-F4, Fp1-F3) were excluded, as these are most susceptible to muscle and eye-blink artifacts. If any of the remaining leads contained artifacts or excessive noise, they were replaced by the best-quality frontal or temporal lead available. For hiPSC cultures on 24-well MEAs (12 electrodes per well), the first and last electrodes were excluded by default, yielding 10 electrodes. If any of these showed artifacts or high noise levels, they were replaced with the first or last electrode, provided it was artifact-free. For primary cultures on 60-electrode single-well MEAs, 10 electrodes were selected to mimic the spatial layout of the 24-well configuration (excluding first and last positions). Artifactual or noisy electrodes from this selection were replaced with nearby artifact-free electrodes.

All signals (EEG, MEA, and simulations) underwent identical preprocessing: zero padding to minimize filter edge effects, high-pass filtering at 0.1 Hz, low-pass filtering at 25 Hz, trimming of filter transients, and downsampling to 250 Hz.

Signals were segmented into 50-second epochs, matching the shortest continuous recording available from healthy subjects. Epochs with visually identified artifacts were excluded. For BS recordings, epochs containing fewer than two bursts were discarded. All epochs were selected from periods previously annotated by EEG experts as containing either physiological *α*-activity or BS.

We calculated two metrics for each epoch: (1) joint Lempel-Ziv complexity for multivariate signals^71^, measuring signal predictability, and (2) burstiness, defined as the coefficient of variation of the absolute derivative of voltage recordings, averaged across channels, quantifying signal intermittency. Both metrics were averaged across all available epochs per subject.

### Computational model and *in silico* experiments

For *in silico* experiments, we used our previously validated computational model of hiPSC-derived neuronal networks^37^. The model comprises 100 Hodgkin-Huxley neurons with voltage-gated sodium and potassium currents plus spike-triggered slow afterhyperpolarizing currents. Neurons were randomly connected via conductance-based synapses containing AMPA and NMDA receptors with short-term synaptic depression. Neurons are heterogeneously excitable and receive independent noisy membrane potential fluctuations, mimicking the effect of membrane or synaptic noise. Ultimately, bursts are initiated by the spontaneous firing of a few highly excitable neurons when their membrane potential fluctuations reach the firing threshold.

We employed simulation-based inference (SBI) to estimate model parameters from experimental MEA data as done previously^72^. We sampled 500,000 parameter configurations from uniform prior distributions and performed 3-minute simulations for each configuration. From these simulations, we extracted 15 MEA features capturing network activity characteristics. A neural density estimator (Masked Autoregressive Flow) was trained on these parameter-feature pairs to learn the mapping between MEA features and model parameters. From our hiPSC-derived neuronal network MEA measurements, we computed the same 15 features and evaluated them using the trained estimator to obtain posterior distributions over model parameters, representing the probability of different parameter combinations given the experimental observations. We used the mode of the posterior distribution as our baseline parameter values for subsequent analyses. Model simulations and simulation-based inference were performed in Python 3.9 using the brian2^73^ and sbi^74^ packages, respectively.

For LZ complexity and burstiness quantification, we took the signal from the virtual electrodes in our computational model and filtered and downsampled identically to how we processed the EEG and MEA signals, with 5-minute simulations used for quantification.

To simulate the effect of inhibition, we added 40 inhibitory neurons (70:30 excitatory:inhibitory ratio) with identical intrinsic properties to excitatory neurons. GABA synapses had -60 mV reversal potential, instantaneous opening, and exponential decay (*τ* = 50 ms)^75^. Maximum GABA conductance was systematically varied until activity features saturated.

To simulate the effect of modularity, we replicated the experimental set-up of Yamamoto et al. *in silico*^36^. We divided the neurons into 4 equally occupied clusters and defined parameter Q, determining the ratio of intra-module connections to inter-module connections. A Q of 1 suggests a completely random network without structural modularity. For higher values of Q, the number of connections within modules becomes larger than between modules, while keeping the overall number of synapses the same.

To simulate the effect of increased input, we increased the noise parameter of our computational model. This parameter governs the standard deviation of the normal distribution from which membrane potential fluctuations are drawn. This noise differs per neuron. These membrane potential fluctuations simulate the response of the membrane to both channel noise and synaptic noise. Synaptic noise would increase if the amount of subcortical inputs increases, and it therefore serves as a proxy for this. Since increased noise dramatically elevated firing rates and burst frequencies, we proportionally reduced neuronal excitability by adjusting input current from +13 pA (no noise) to -10 pA (highest noise), proportional to the noise level, to maintain comparable activity levels.

### Statistical analysis

Statistical analyses were performed using GraphPad Prism 5 (GraphPad Software, Inc., CA, USA). After assessing normality with Kolmogorov-Smirnov tests, we used Wilcoxon matched-pairs signed rank tests for paired comparisons, Kruskal-Wallis tests with Dunn’s multiple comparisons for group comparisons, and two-way ANOVA with Dunnett’s multiple comparisons for network randomness experiments. Significance was set at *p <* 0.05.

## Data and Code availability

EEG data, MEA data, and simulations are available on figshare (doi: 10.6084/m9.figshare.29859185). All MATLAB code to perform data analysis and Python code to perform simulations and fit parameters can be found on GitLab (https://gitlab.utwente.nl/m7706783/burst-suppression).

## Acknowledgements

We thank Hideaki Yamamoto and Jordi Soriano-Fradera for kindly and promptly providing the experimental data of their modular cultures. We also thank Sönke van Loh for his helpful guidance on complexity measures and their applications. Furthermore, we thank Marloes Levers for her help with the *in vitro* culturing and recording of hiPSC-derived neuronal networks.

## Author Contributions

N.D. conducted data analysis and visualization and wrote the manuscript with input from all authors. N.D. and G.H. performed *in vitro* experiments. N.D. and M.vP. conceptualized the project. M.vP. curated the EEG data. M.F. acquired funding. M.F. and M.vP. supervised the project.

