## Supplementary Figures for "Burst suppression: a default brain state associated with loss of network complexity"

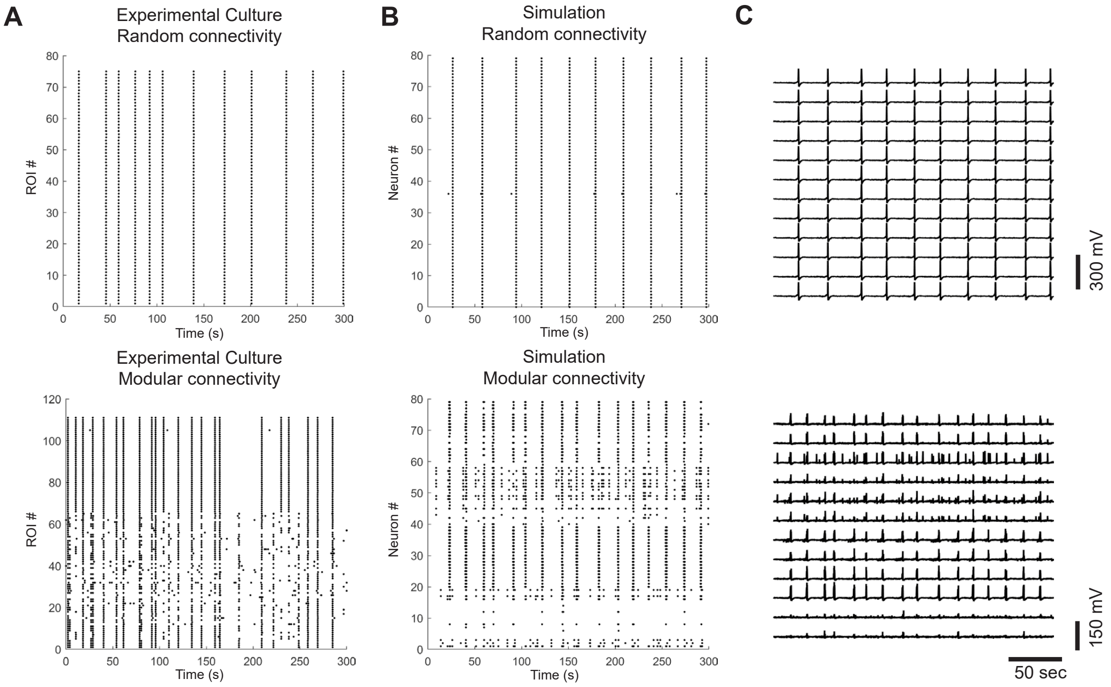


Figure S1: **Simulations reproduce experimental results of neuronal cultures with modular architecture by Yamamoto et al.**
**A)** Raster plots of detected calcium events from primary cultures, data kindly provided by Hideaki Yamamoto and originally reported in[1]. Top shows activity of a merged randomly connected network of cultured neurons. Bottom shows activity of a network with four sparsely connected modules of neurons. **B)** Rasterplots of simulations with our computational model, note these represent action potentials while the experimental results show calcium events (bursts of action potentials). Top shows simulations with a randomly connected merged model, and bottom shows simulations with a model where neurons are divided into four equally occupied modules that are sparsely connected. **C)** Signals from the virtual electrodes of the simulations shown in panel B, filtered in the EEG band.

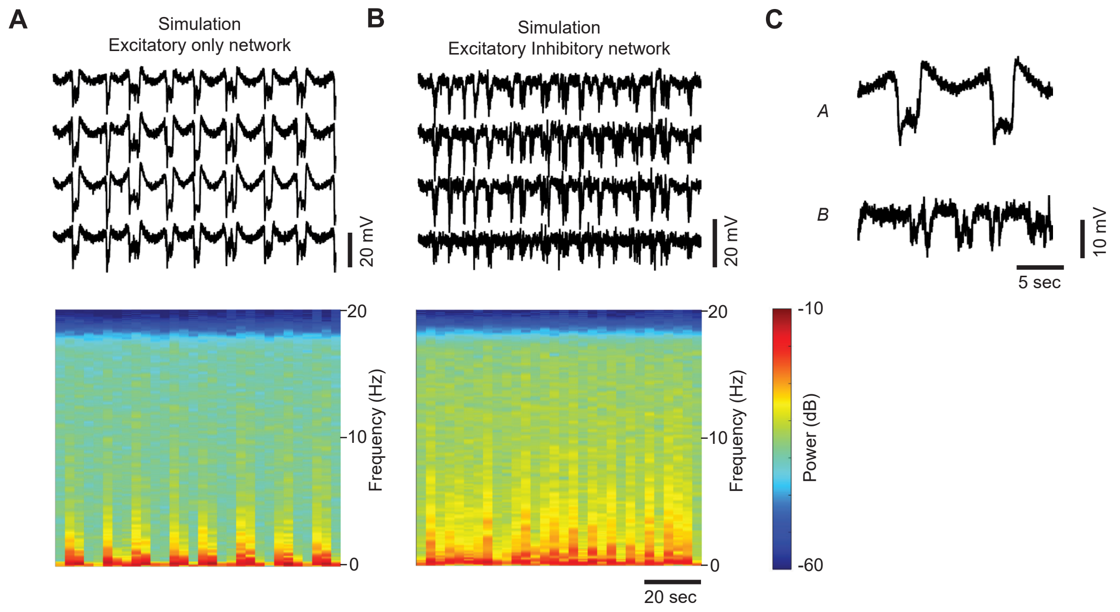


Figure S2: **Frequency content and voltage zooms of neuronal network simulations** **A-B)** Top: 100 s of virtual MEA signal (filtered 0.1-25 Hz) from four representative electrodes and bottom: average spectrogram from all virtual electrodes from **A)** simulations with the neuronal network model fitted to hiPSC-derived neuronal network recordings (note the similarity to Figure 1), and **B)** simulations with the neuronal network model after the addition of inhibitory neurons. **C)** Representative 20 s epochs from one electrode of the respective panels A and B. Note the clear difference between identical bursts (top) and heterogeneous bursts (bottom).
